# Hippocampal knockdown of Piwil1 and Piwil2 enhances contextual fear memory in mice

**DOI:** 10.1101/298570

**Authors:** Laura J. Leighton, Wei Wei, Vikram Singh Ratnu, Xiang Li, Esmi L. Zajaczkowski, Paola A. Spadaro, Nitin Khandelwal, Arvind Kumar, Timothy W. Bredy

**Affiliations:** Cognitive Neuroepigenetics Laboratory, Queensland Brain Institute, The University of Queensland, Brisbane, QLD, 4072, Australia.; CSIR-Centre for Cellular and Molecular Biology (CCMB), Hyderabad, 500007, India.

## Abstract

The Piwi pathway is a conserved gene regulatory mechanism comprised of Piwi-like proteins and Piwi-interacting RNAs, which modulates gene expression via RNA interference and epigenetic mechanisms. The mammalian Piwi pathway has been defined by its role in transposon control during spermatogenesis, and despite an increasing number of studies demonstrating its expression in the nervous system, relatively little is known about its function in neurons or potential contribution to gene regulation in the brain. We have discovered that all three Piwi-like genes are expressed in several regions of the mouse brain, and that simultaneous knockdown of *Piwil1* and *Piwil2* in the adult mouse hippocampus enhances contextual fear memory without affecting generalised anxiety. Our results implicate the Piwi pathway in control of plasticity-related gene expression in the adult mammalian brain.

## Introduction

Learning-related neuronal plasticity and the formation of memory depend on tightly controlled patterns of gene expression, with non-coding RNAs (ncRNAs) emerging as an essential regulatory mechanism in this process (Bredy, Lin, Wei, Baker-Andresen, and Mattick, 2011; Leighton, Ke, Zajaczkowski, Edmunds, Spitale, and Bredy, 2017; Spadaro and Bredy, 2012). Non-coding RNAs are diverse in size, structure, origin and function; small ncRNAs, in particular, arise through a number of distinct biogenesis pathways, are categorised into several distinct classes, and primarily serve to fine-tune gene expression in response to current environmental demands. The best characterised small ncRNAs are microRNAs (miRNAs), which have known roles in neuronal function, cognition, learning and memory (reviewed by Fiorenza and Barco, 2016). For instance, the brain-enriched miR-128b is involved in regulating fear extinction memory (Lin, Wei, Coelho, Li, Baker-Andresen, Dudley, Ratnu, Boskovic, Kobor, and Sun, 2011), and learning-induced downregulation of miR-182 supports fear memory formation (Griggs, Young, Rumbaugh, and Miller, 2013). Although these and other studies (reviewed by Bredy et al., 2011) have focused on the importance of neuronal miRNAs, there are many other classes of small ncRNAs that are expressed in the brain but have received relatively little attention in the context of neuronal function and learning.

One such class of small ncRNAs are the Piwi-interacting RNAs (piRNAs), which together with the Piwi-like proteins (a subset of the Argonaute family) form an evolutionarily-conserved gene regulatory mechanism often referred to as the Piwi pathway. Characteristic features of piRNAs include a length of 26-31nt, occurrence in genomic clusters, a bias towards uracil as the first base, and 2’-O-methylation of the terminal nucleotide (Girard, Sachidanandam, Hannon, and Carmell, 2006; Grivna, Beyret, Wang, and Lin, 2006; Kirino and Mourelatos, 2007; Ohara, Sakaguchi, Suzuki, Ueda, Miyauchi, and Suzuki, 2007); however, there is some variability between species in the size, function and cell-type specificity of piRNAs. In mammals, Piwi proteins and piRNAs are found primarily within the male germline, where they are necessary for germ cell maintenance and spermatogenesis because they protect the genome by silencing transposon expression at both the epigenetic and post-transcriptional levels (Carmell, Girard, van de Kant, Bourc’his, Bestor, de Rooij, and Hannon, 2007; Deng and Lin, 2002; Iwasaki, Siomi, and Siomi, 2015; Kuramochi-Miyagawa, Kimura, Ijiri, Isobe, Asada, Fujita, Ikawa, Iwai, Okabe, and Deng, 2004). In the mouse testis, genomic piRNA clusters are transcribed and processed into individual piRNAs via several processing steps which are incompletely understood; in addition, Piwi proteins guided by these primary piRNAs cleave complementary transposon transcripts, which are then processed into secondary piRNAs in an amplification loop referred to as the ‘ping pong cycle’ (reviewed by Iwasaki et al., 2015).

In addition to their role in germline genome defence, there is a growing recognition that the Piwi pathway is involved in cell fate decisions during development and throughout the lifespan (Ponnusamy, Yan, Liu, Li, and Wang, 2017). This function extends to neuronal precursors; *Piwil1* is involved in the polarisation and radial migration of neurons in the mouse fetal cerebral cortex, partly through its effect on the expression of microtubule-associated proteins (Zhao, Yao, Chang, Gou, Liu, Qiu, and Yuan, 2015), and the neocortical circuitry is disrupted in the *Piwil1* knock-out mouse model, compromising development (Viljetic, Diao, Liu, Krsnik, Wijeratne, Kristopovich, Dutre-Clarke, Kraushar, Song, and Xing, 2017). These findings are consistent with the strong association between coding mutations of *Piwi* genes in humans, and autism (Iossifov, O’roak, Sanders, Ronemus, Krumm, Levy, Stessman, Witherspoon, Vives, and Patterson, 2014).

The Piwi pathway is also involved in neuronal gene regulation in the adult brain. In 2012, Rajasethupathy *et al*. reported that *Aplysia californica* expresses a single Piwi-like protein and dozens of piRNAs within neurons (Rajasethupathy, Antonov, Sheridan, Frey, Sander, Tuschl, and Kandel, 2012); knockdown of *PIWI* or one specific piRNA is sufficient to impair long-term facilitation, which is a form of neural plasticity thought to underlie learning and memory. Several other studies suggest that the Piwi pathway reprises this role in mammalian neurons. Lee *et al*. detected PIWIL1 in mouse cortical neurons *in vitro* (Lee, Banerjee, Zhou, Jammalamadaka, Arcila, Manjunath, and Kosik, 2011) and also reported several dozen putative piRNAs, although a recent reanalysis has shown that they are unlikely to be *bona fide* piRNAs (Tosar, Rovira, and Cayota, 2018). A more recent exploration of neuronal piRNAs identified approximately 30,000 piRNAs from mouse brain tissue (Ghosheh, Seridi, Ryu, Takahashi, Orlando, Carninci, and Ravasi, 2016) and suggested that these brain-derived piRNAs are most similar to those associated with PIWIL2 in mouse testes. Another study identified low-level expression of truncated PIWIL2 in the adult mouse brain, with hypomethylation of transposons and behavioural defects observed in *Piwil2* knockout mice (Nandi, Chandramohan, Fioriti, Melnick, Hébert, Mason, Rajasethupathy, and Kandel, 2016). We have also previously demonstrated that the Piwi-like genes *Piwil1* and *Piwil2* are expressed in cultured mouse neurons, and have shown that they are dynamically regulated by neuronal activation, and are involved in transcriptional regulation of activity-induced gene expression (Leighton, Zhao, Li, Dai, Marshall, Liu, Wang, Zajaczkowski, Khandelwal, and Kumar, 2018).

Here we demonstrate that all three murine Piwi proteins are expressed in the adult mouse brain, and that a functional disruption of the two most abundant, namely *Piwil1* and *Piwil2*, in the hippocampus produces a striking enhancement of contextual fear memory. These findings suggest that, together with other classes of small ncRNAs, the Piwi pathway is also involved in learning and memory.

## Materials and methods

### Animals

Male C57BL/6 mice (14 weeks old) were housed in divided cages on a 12-hour light cycle with food and water available *ad libitum*. Behavioural testing was conducted during the light phase. All protocols involving animals were approved by the Animal Ethics Committee of The University of Queensland. Animal experiments were carried out in accordance with the Australian Code for the Care and Use of Animals for Scientific Purposes (8^th^ edition, revised 2013.)

### Adeno-associated virus construction and packaging

Short hairpin RNAs (shRNAs; one targeting each of *Piwil1* and *Piwil2*, and a non-targeting control) were designed, cloned into the pAAV2-mu6 vector, and packaged into adeno-associated viruses (AAVs) as described previously (Leighton et al., 2018). Sequences of shRNAs were previously published (Leighton et al., 2018).

### Culture of primary mouse cortical neurons

Primary neuronal cultures were prepared from E16 mouse cortices as described previously (Leighton et al., 2018). To knock down expression of genes of interest, cultured neurons at 2-3 days *in vitro* were exposed to AAV for 12 hours, followed by replacement of the culture medium. Cells were collected for molecular analysis 7 days after treatment with AAVs.

### RNA isolation and reverse transcription

To extract RNA, cells or tissue were lysed with Nucleozol (Macherey-Nagel) and RNA was isolated following the manufacturer’s instructions. Reverse transcription was carried out using the QuantiTect kit (QIAGEN) using the provided RT Primer Mix and following the manufacturer’s instructions.

### Quantitative PCR

Quantitative PCR reactions were prepared in duplicate, in a 10µL reaction volume, using 2X SYBR master mix (QIAGEN), 500µM of each primer and 1µL per reaction of a cDNA sample (the dilution of which varied according to target abundance). Reactions were run on the Rotor-Gene Q platform and results analysed using the delta-delta-CT method, normalised to the reference gene *Pgk1* (phosphoglycerate kinase). Primer sequences were previously published (Leighton et al., 2018).

### Intrahippocampal injection of AAV vector

Male C57BL/6 mice (14 weeks old) were anaesthetised by interperitoneal injection of 100mg/kg ketamine and 10mg/kg xylazine, and placed in a stereotaxic frame. The skull was exposed and burr holes drilled at positions of interest. AAV vector (~1.3µL per side) was injected into the hippocampus at coordinates AP -2.46, ML ± 1.7 DV -1.8 relative to bregma, using a pulled glass micropipette. Needles were withdrawn after 10 minutes and the skin incision closed with surgical glue. Mice were allowed to recover for 2 weeks before behavioral experiments.

### Contextual fear conditioning and recall testing

Contextual fear conditioning was performed in individual soundproof operant chambers with plastic walls and a wire grid floor, which were scented with lemon essence and cleaned with hot water and ethanol in between animals. Mice spent a total of 8 minutes in the apparatus; 1-second, 0.7mA footshocks were delivered at 2, 4 and 6 minutes. Recall tests were performed 24 hours and 7 days after fear conditioning, by returning the mice to the apparatus for 2 minutes. Data was analysed using FreezeFrame 4 (Actimetrics) by a researcher blind to treatment group of the mice. Statistical analysis was performed using a two-way mixed ANOVA with time as the repeated-measures factor and experimental group as the between-subjects factor. Groups were compared within each time point using a Bonferroni multiple comparisons correction.

### Elevated plus maze

Elevated plus maze testing was conducted in a sound-attenuated behavioral room moderately illuminated with white light at 150 ± 3 lux. The elevated plus maze was constructed of grey acrylic. Each arm was 30cm long and 5cm wide, with the centre zone measuring 5×5 cm, and the arms were 40cm off the floor. Mice were gently placed in the centre zone of the maze and recorded for 5 minutes. The maze was cleaned with 70% ethanol and allowed to dry in between mice. Videos were analysed using EthoVision XT 11.5 (Noldus) to determine the percentage of time each mouse spent in the open arms, closed arms and centre zone (judged by the position of the body centrepoint.) Statistical analysis was performed using Student’s t-test with Welch’s correction.

### Open field test

The open field test was conducted in a sound-attenuated behavioral room dimly illuminated with white light at 60 ± 3 lux. Four white plastic open fields (30 x 30 x 30cm) were used; the centre zone was defined as a 15 x 15 cm square concentric with the base of the arena. One mouse was placed gently in the centre of each field and mice were recorded for 20 minutes. Fields were cleaned with 70% ethanol and allowed to dry in between mice. Videos were analysed using EthoVision XT 11.5 (Noldus) to determine the total distance travelled and time spent in the centre zone (judged by the position of the body centrepoint.) Statistical analysis was performed using Student’s t-test with Welch’s correction.

### Histology and imaging

After the completion of behavioural testing (approximately 8 weeks after AAV vector injection), mice were transcardially perfused with 50mL of cold phosphate buffered saline (PBS) followed by 50mL of 4% paraformaldehyde in PBS. Brains were removed and post-fixed in 4% paraformaldehyde overnight at room temperature, then stored in PBS at 4C until processing. Tissue was sectioned at 40 microns using a vibrating microtome. Selected tissue sections were blocked with 5% goat serum and 0.1% Triton X-100 in PBS, then incubated overnight with primary antibody (rabbit polyclonal against GFP, Abcam ab6556, 1:500). After 3x 20 minute washes with PBS, sections were incubated with secondary antibody (Jackson ImmunoResearch, DyLight 549 Goat anti-Rabbit, 1:1000) and DAPI (1:2000), then mounted onto Superfrost slides and coverslipped with DAKO fluorescence mounting medium. Imaging was performed at the Queensland Brain Institute’s Advanced Microscopy Facility, using a Zeiss Axio Imager microscope.

## Results

### Piwi-like genes are expressed in the adult mouse brain

To begin to investigate the function of the Piwi pathway in learning and memory, we first asked whether Piwi-like proteins were present in various regions of the adult mouse brain. Quantitative reverse transcription PCR was used to detect each of the three mouse Piwi genes: *Piwil1* (Miwi), *Piwil2* (Mili) and *Piwil4* (Miwi2). We found that all three Piwi-like genes were expressed in each of the regions of the adult mouse brain examined (prefrontal cortex, hippocampus, somatosensory cortex, and cerebellum) as well as in whole fetal brain at E17 (Figure 1).

**Figure 1:**
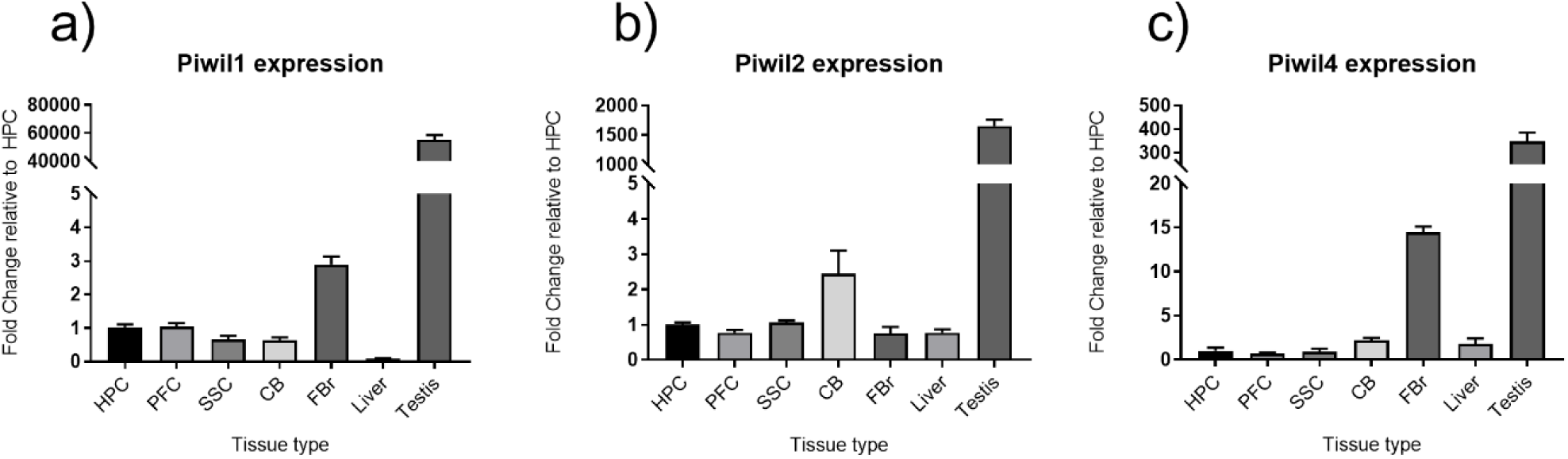
Piwi-like genes are expressed in the adult mouse brain. Expression of *Piwil1* (a), *Piwil2* (b) and *Piwil4* (c) was examined using qRT-PCR in four regions of the mouse brain: hippocampus (HPC), prefrontal cortex (PFC), somatosensory cortex (SSC) and cerebellum (CB), as well as in whole fetal brain (FBr), and adult liver and testis (n=4 for each tissue type).

### Knockdown of Piwil1 and Piwil2

In mice, *Piwil1* and *Piwil2* are involved in primary processing of piRNAs from their precursor transcripts, whereas *Piwil4* is not absolutely required for primary piRNA expression or piRNA amplification (De Fazio, Bartonicek, Di Giacomo, Abreu-Goodger, Sankar, Funaya, Antony, Moreira, Enright, and O’Carroll, 2011; Manakov, Pezic, Marinov, Pastor, Sachidanandam, and Aravin, 2015). Therefore, disruption of *Piwil1* and *Piwil2* would be expected to effectively disable the function of the entire Piwi pathway in mouse neurons. We created AAV vectors carrying shRNAs directed against *Piwil1* and *Piwil2*, and a control shRNA which has no specificity to any known mouse transcripts. We validated the AAV vectors by confirming that the shRNAs against *Piwil1* and *Piwil2* knocked down their respective target mRNAs in cultured primary neurons (Figure 2 a, b).

**Figure 2:**
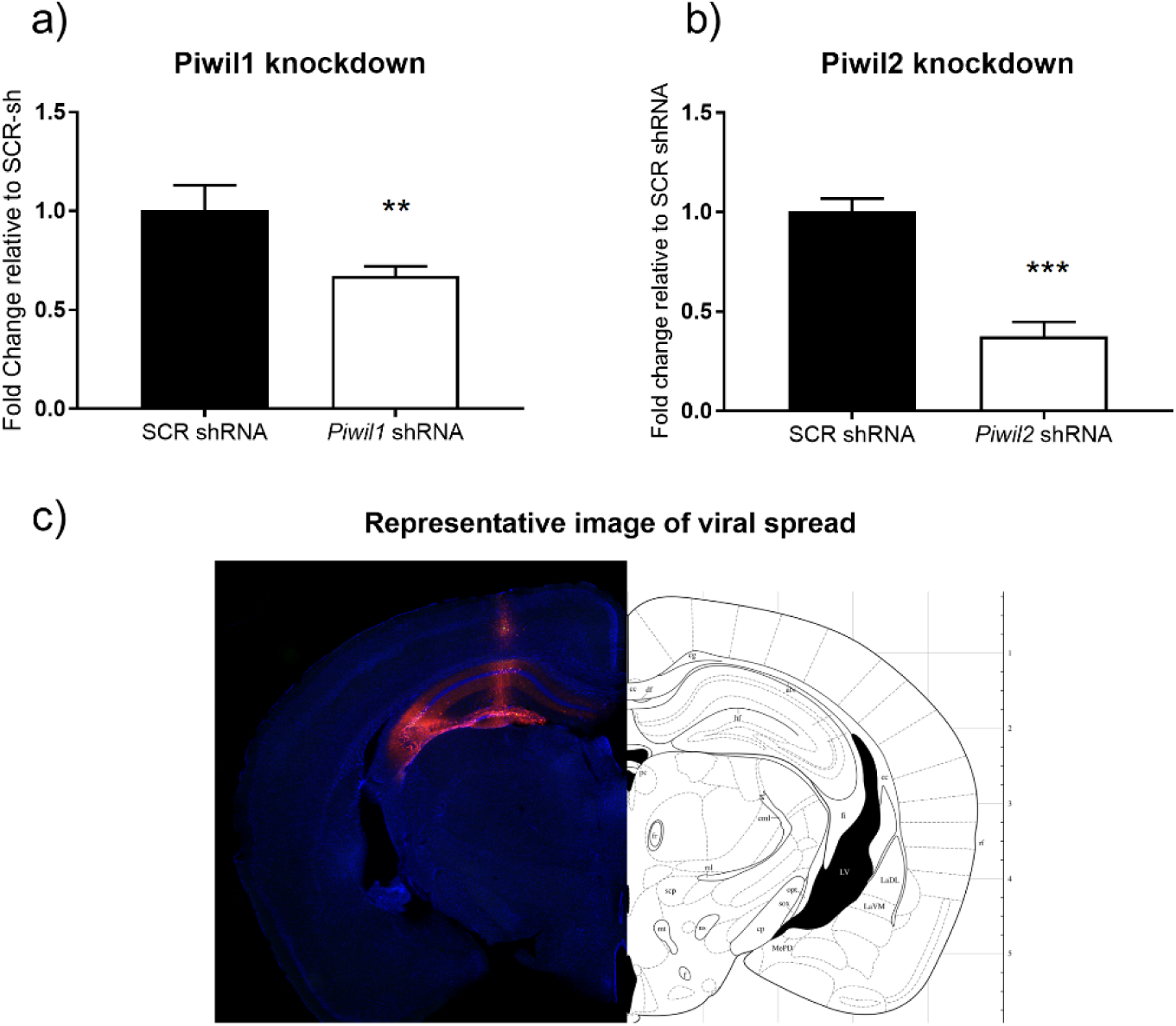
Validation of *Piwil1* and *Piwil2* knockdown. AAV vectors expressing short hairpin RNAs targeting *Piwil1* (a) and *Piwil2* (b) produce significant knockdown of their respective targets in cultured mouse neurons, relative to the non-targeting control (*Piwil1* knockdown n=4, Student’s t-test, p=0.0082; *Piwil2* knockdown n=4, Student’s t-test, p=0.0006) Representative images of viral spread within the adult mouse hippocampus (g) show effective targeting of the hippocampus by the AAV.

### Hippocampal injection of shRNA-expressing AAV vectors

To examine the function of the Piwi pathway in hippocampal-dependent learning, we bilaterally injected AAV vectors into the hippocampi of C57BL/6 mice. Animals in the Piwi-sh group received AAVs expressing shRNAs targeting both *Piwil1* and *Piwil2* (mixed together in a 1:1 ratio) while animals in the control group received AAV expressing control shRNA. The coordinates used for injection reliably resulted in diffuse spread of the vector through the hippocampus, with minimal infection of overlying cortical structures (Figure 2c).

### Disruption of the Piwi pathway enhances contextual fear memory

Following a 2-week post-surgery recovery period, mice from the Piwi-sh group (n=10) and scrambled-control group (n=11) underwent contextual fear conditioning consisting of an 8-minute exposure to a novel environment, with 1-second, 0.7mA footshocks delivered at 2, 4 and 6 minutes. Mice from both groups showed normal acquisition of conditioned fear (Figure 3a). However, there was significantly greater freezing in Piwi-sh mice relative to control mice during recall tests both 24 hours and 7 days after fear conditioning (Figure 3b), revealing an enhancement of long-term memory.

**Figure 3:**
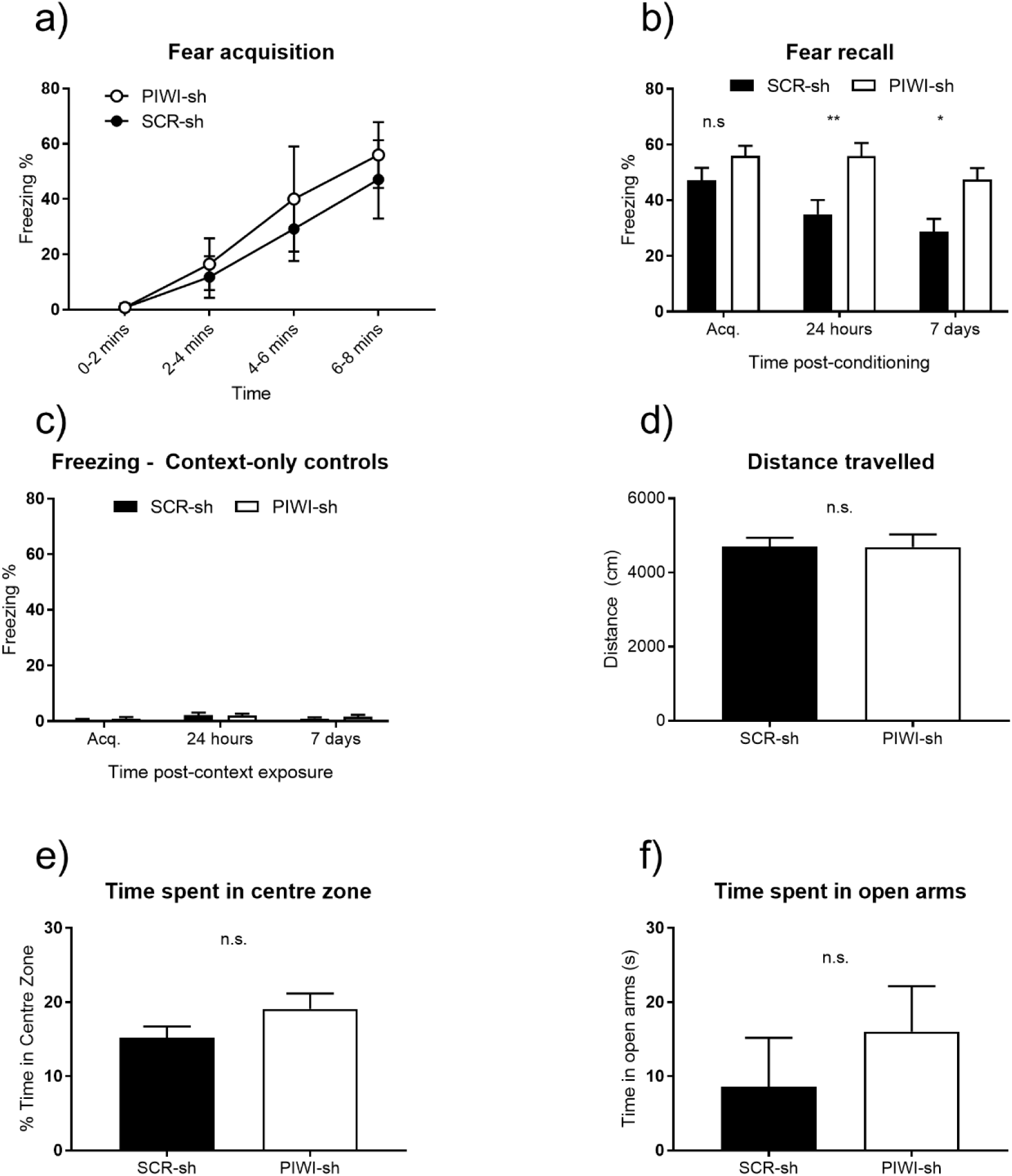
Knockdown of *Piwil1* and *Piwil2* enhances contextual fear memory. Animals injected with AAVs targeting both *Piwil1* and *Piwil2* (‘PIWI-sh’) show normal acquisition of contextual fear (a) compared to animals injected with non-targeting control AAV (‘SCR-sh’) (two-way ANOVA, t=1.401, p_adj._=0.5001.) However, PIWI-sh animals show increased freezing (b), corresponding to enhanced memory, 24 hours (t=3.335, p_adj._=0.0045) and 7 days (t=2.967, p_adj._=0.0132) after conditioning. Animals from either group exposed to the training context but never shocked (c) show no fear. There is no difference between PIWI-sh and SCR-sh animals in distance travelled (d; Student’s t-test, p=0.9791) or time spent in the centre zone (e; Student’s t-test, p=0.1432) during a 20-minute open field test; there is also no significant difference in time spent in the open arms of the elevated plus maze (f; Student’s t-test, p=0.2885)

### Disruption of the Piwi pathway does not affect anxiety or locomotion

Given that we observed an increase in freezing behaviour in Piwi-sh mice compared to controls, we asked whether these animals showed any alteration in generalised anxiety or locomotion that could offer an alternative explanation for this effect. We found no difference in distance travelled between the Piwi-sh (n=11) and control mice (n=10) during a 20 minute open field trial (Figure 3d), indicating no effect of Piwi knockdown on locomotion or exploratory behaviour. We also found that there was no difference between groups in the time spent in the centre of the open field during a 20 minute trial (Figure 3e) or in the open arms of the elevated plus maze during a 5 minute trial (Figure 3f). Together with the observation that Piwi-sh mice exposed to the fear conditioning context but never shocked are not afraid (Figure 3c), these results show that disruption of the hippocampal Piwi pathway has no effect on generalised or novelty-responsive anxiety.

## Discussion

Despite several reports of piRNA expression in the mammalian brain, including piRNA dysregulation in response to manipulation or disease, very little research has focused on the functional role of Piwi proteins or the Piwi pathway as a whole. In this study, we have demonstrated that disruption of this pathway via knockdown of *Piwil1* and *Piwil2* produces a robust enhancement of contextual fear memory which is independent of related or confounding behaviours, including locomotion and anxiety. Our results provide strong justification for further exploration of the Piwi pathway as a neuroepigenetic regulator of experience-dependent gene expression in the adult brain.

Several recent studies have implicated the Piwi pathway in brain function through neurodevelopmental phenotypes. Zhao *et al*. report that *Piwil1* regulates polarity and migration of neurons in the developing mouse neocortex (Zhao et al., 2015), and a *Piwil1* knockout mouse strain has been shown to exhibit disrupted neocortical development and circuitry (Viljetic et al., 2017). These reports are congruent with the finding that *de novo* coding mutations in human *Piwil* genes are strongly associated with autism-spectrum disorders (Iossifov et al., 2014) which are increasingly understood to involve aberrant organisation of cortical neurons (Nunes, Peatfield, Vakorin, and Doesburg, 2018; Packer, 2016). In addition, Nandi *et al*. found that *Piwil2* knockout mice are hyperactive and that males show reduced anxiety (Nandi et al., 2016); because the genetic absence of *Piwil2* is present from conception, it is likely that this phenotype is also developmental. In the present study, we have demonstrated for the first time that targeted manipulation of the Piwi pathway in the adult mammalian brain affects behaviour. Strikingly, the behavioural alteration that we describe is a gain of function rather than an impairment. This strongly suggests that the Piwi pathway is functioning as an adaptive and dynamic mechanism for gene regulation in post-mitotic neurons of the mouse hippocampus. This finding complements the demonstration of rapid, adaptive gene regulation that is modulated by the Piwi pathway in cultured neurons of *Aplysia* (Rajasethupathy *et al*., 2012).

A number of studies have detected the expression of piRNAs in the mammalian brain (Dharap, Nakka, and Vemuganti, 2011; Ghosheh et al., 2016; Lee et al., 2011; Nandi et al., 2016; Qiu, Guo, Lin, Yang, Zhang, Zhang, Zuo, Zhu, Li, and Ma, 2017; Roy, Sarkar, Parida, Ghosh, and Mallick, 2017; Saxena, Tang, and Carninci, 2012). The criteria used to classify small RNAs such as piRNAs have varied, and in some cases, most of the putative piRNAs reported are actually fragments of other RNA species including ribosomal RNA, transfer RNA and small nucleolar RNA (Tosar et al., 2018). However, two recent studies used stringent criteria to define putative piRNAs, and identified a population of small RNAs from the mouse brain with virtually all the hallmarks of piRNAs, including size, 1U-bias, and genomic organisation within known piRNA clusters (Ghosheh et al., 2016; Nandi et al., 2016). Both of these studies reported that the majority of brain-derived piRNAs are directed against transposons, particularly LINE1 elements. Transposon activity, particularly of LINE1 elements, is a recognised mechanism for generating functionally relevant somatic mosaicism within hippocampal neurons that is associated with learning and memory (Bachiller, del-Pozo-Martín, and Carrión, 2017; Baillie, Barnett, Upton, Gerhardt, Richmond, De Sapio, Brennan, Rizzu, Smith, and Fell, 2011; Kurnosov, Ustyugova, Nazarov, Minervina, Komkov, Shugay, Pogorelyy, Khodosevich, Mamedov, and Lebedev, 2015), and regulation of this process is a probable mechanism for the influence of the Piwi pathway on learning and memory. This hypothesis is supported by recent findings that overexpression of *Piwil1* protects neurons *in vitro* and *in vivo* from pathologic LINE1 derepression and DNA breakage caused by oxidative stress or genetic insufficiency of other transposon control pathways (Rekaik, Peze-Heidsieck, Massiani-Beaudoin, Joshi, Fuchs, and Prochiantz, 2017). Another potential mechanism for hippocampal piRNA function is offered by the presence of the hippocampal neurogenic niche; given the broad role of the Piwi pathway in maintenance of stemness and asymmetric division of stem cells, it is possible that Piwi activity is involved in regulation of hippocampal neurogenesis. Interestingly, the behavioural phenotype that we observe in this study (increased freezing to a conditioned stimulus 24 hours and 7 days post-training) may be interpreted as either ‘enhanced memory’ or ‘attenuated forgetting’. Hippocampal neurogenesis is associated with forgetting because integration of newborn neurons into existing hippocampal circuitry reduces expression of existing memories (Akers, Martinez-Canabal, Restivo, Yiu, De Cristofaro, Hsiang, Wheeler, Guskjolen, Niibori, and Shoji, 2014; Gao, Xia, Guskjolen, Ramsaran, Santoro, Josselyn, and Frankland, 2018); it is plausible that disruption of the Piwi pathway may impair neurogenesis by causing dysfunction of stem cells or transient neuronal progenitors. This is supported by the finding that *Piwil1* and *Piwil2* are upregulated in differentiating neurospheres compared to both proliferating and differentiated cells (N. Khandelwal, personal communication.) However, detection of Piwi proteins and piRNAs in regions of the adult brain without neurogenic stem cells implies that they also have functions in post-mitotic neurons.

Finally, earlier work by our laboratory determined that *Piwil1* and *Piwil2* are dynamically upregulated by stimulation of cultured mouse neurons, and that they promote activity-dependent chromatin relaxation at the calcineurin (*Ppp3r1)* locus which is required for binding of the regulatory factor ING1 and subsequent expression of *Ppp3r1* (Leighton *et al*., 2018). In that study, we proposed that the Piwi pathway, which is known to respond to DNA damage (Yin, Wang, Chen, Liu, Han, Yan, Shen, He, Duan, and Li, 2011), may target this and other loci by recognising DNA double-strand breaks (DSBs) which are a feature of rapid adaptive transcription in neurons (Madabhushi, Gao, Pfenning, Pan, Yamakawa, Seo, Rueda, Phan, Yamakawa, and Pao, 2015). This is an attractive explanation for neuronal Piwi function because the mechanism is potentially applicable to any of the dozen or more genes whose transcription is facilitated by DSBs. However, additional work is necessary to determine whether DSBs recruit Piwi proteins, and whether other DSB-regulated genes are also Piwi targets.

In summary, the nature and function of the Piwi pathway in neurons remains enigmatic. Here, we have added to an emerging body of evidence that this conserved gene regulatory mechanism is functionally relevant within the adult hippocampus. Our finding that a disruption of the Piwi pathway in neurons results in memory enhancement strongly implies its involvement in rapid and adaptive transcriptional or post-transcriptional regulatory processes. Further study is needed to identify the cell-type specificity and regulatory targets of the Piwi pathway in order to understand its contribution to neuronal gene regulation.

## Acknowledgements

The authors gratefully acknowledge grant support from the NIH (1R21MH103812 AND 5R01MH105398-TWB) and the NHMRC (APP1069570 AND APP1083468-TWB). L.J.L. is supported by postgraduate scholarships from the Westpac Bicentennial Foundation and the Australian Government Research Training Program. We would also like to thank Ms. Rowan Tweedale for helpful editing of the manuscript.

